# RoboEM: automated 3D flight tracing for synaptic-resolution connectomics

**DOI:** 10.1101/2022.09.08.507122

**Authors:** Martin Schmidt, Alessandro Motta, Meike Sievers, Moritz Helmstaedter

## Abstract

Mapping neuronal networks from 3-dimensional electron microscopy data still poses substantial reconstruction challenges, in particular for thin axons. Currently available automated image segmentation methods, while substantially progressed, still require human proof reading for many types of connectomic analyses. RoboEM, an AI-based self-steering 3D flight system trained to navigate along neurites using only EM data as input, substantially improves automated state-of-the-art segmentations and replaces human proof reading for more complex connectomic analysis problems, yielding computational annotation cost for cortical connectomes about 400-fold lower than the cost of manual error correction.

---

Extracting the dense neuronal connectivity from 3D EM data of brain tissues poses major computational challenges (Helmstaedter, Briggman et al. 2013, Kasthuri, Hayworth et al. 2015, Wanner, Genoud et al. 2016, Eichler, Li et al. 2017, Motta, Berning et al. 2019, Scheffer, Xu et al. 2020). Substantial progress in the field has allowed us to move from fully manual skeleton reconstructions of neurites (White, Southgate et al. 1986, Bock, Lee et al. 2011, Helmstaedter, Briggman et al. 2011, Lee, Bonin et al. 2016, Wanner, Genoud et al. 2016, Eichler, Li et al. 2017, Schmidt, Gour et al. 2017) via combinations of skeleton reconstruction and automated segmentations (Helmstaedter, Briggman et al. 2013, Kornfeld, Januszewski et al. 2020) to proofreading of automated image segmentation (Fig. 1a). This proofreading was initially as laborious as fully manual skeleton-reconstructions (Mishchenko, Hu et al. 2010, Takemura, Bharioke et al. 2013, Kim, Greene et al. 2014, Kasthuri, Hayworth et al. 2015) but has recently been made substantially more efficient by focused human intervention based on much improved automated segmentations (Motta, Berning et al. 2019, Dorkenwald, McKellar et al. 2020, Kornfeld, Januszewski et al. 2020, Xu, Januszewski et al. 2020). Yet, even for automated methods claiming super-human performance (Lee, Zung et al. 2017) or full automation (Kornfeld, Januszewski et al. 2020), when applied to real-world large-scale EM datasets, massive manual annotation efforts are required to obtain scientifically useable connectomes (Zheng, Lauritzen et al. 2018, Scheffer, Xu et al. 2020, MICrONS Consortium, Bodor et al. 2021, Shapson-Coe, Januszewski et al. 2021).

**Figure 1.**
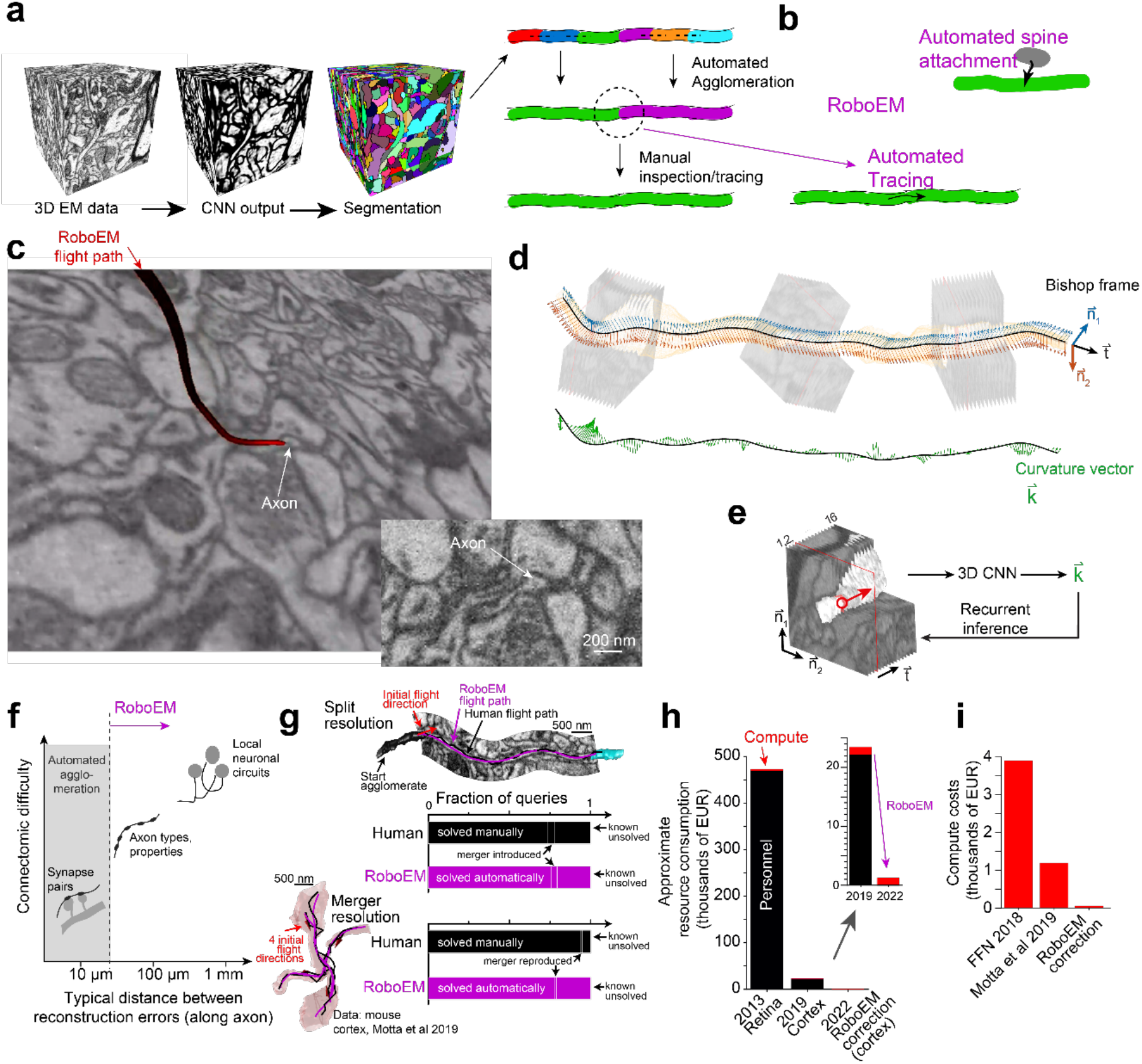
Automated neurite tracing for substitution of human annotation needs in neuronal network reconstruction. (a) Current standard analysis workflow for connectomic analyses from 3D EM data: an initial volume segmentation is generated via CNN-based voxel-wise image analysis (either in two-step process of classification and watershed (Jain, Murray et al. 2007, Turaga, Briggman et al. 2009, Turaga, Murray et al. 2010, Berning, Boergens et al. 2015, Lee, Zung et al. 2017, Funke, Tschopp et al. 2019, Sheridan, Nguyen et al. 2021), or directly as foreground definition, (Meirovitch, Matveev et al. 2016, Januszewski, Kornfeld et al. 2018, Meirovitch, Mi et al. 2019)). Then, based on properties of the interface between segments, an automated agglomeration is generated that aims to combine segmentation objects into longer stretches of axons. Subsequently, all current segmentation methods require manual inspection to resolve remaining errors and reach reconstruction quality that can be used for meaningful connectomic analysis (Plaza 2016, Haehn, Kaynig et al. 2018, Zhao, Olbris et al. 2018, Dorkenwald, Turner et al. 2019, Motta, Berning et al. 2019, Dorkenwald, McKellar et al. 2020, Hubbard, Berg et al. 2020, Kornfeld, Januszewski et al. 2020, Scheffer, Xu et al. 2020, MICrONS Consortium, Bodor et al. 2021). **(b)** RoboEM was built to replace the human inspection and correction step by an automated connection/validation flight that should solve split errors, attach remaining spine heads, and also replace human annotation for merger resolution (Motta, Berning et al. 2019). **(c)** Example of RoboEM flight path along thin axon in SBEM data (Motta, Berning et al. 2019). **(d**,**e)** Design of RoboEM: based on volumetric input data, a steering vector is predicted that, after application, yields novel input data to be processed. Due to its rotation-minimizing property the Bishop frame is a convenient framework to describe position and orientation of the input for RoboEM, and associated Bishop curvatures were used as the steering vector to be predicted. During training, a segmentation mask was used to ‘teach’ corrective steering signals from off-center locations (see Supplementary Figure 1b). **(f)** Calibration of connectomic reconstruction automation by the difficulty of connectomic analyses. Analyses based on synaptic pairs can often be performed already with correct path length of about 10μm, which has already been automatically achieved (Dorkenwald, Turner et al. 2019, Motta, Berning et al. 2019, Kornfeld, Januszewski et al. 2020). For extraction of axonal properties, stretches of typically about 5 synapses are usually required, placing stronger requirements on reconstruction accuracy, see Fig. 2. **(g)** RoboEM performance in direct comparison to human annotators on ending and chiasma queries for split and merge error resolution from (Motta, Berning et al. 2019) **(h)** Effect on resource consumption for connectomic dense reconstructions. Personnel costs are largely removed, focusing on computational costs and the need for increased computational efficiency. Exemplary compute costs for state-of-the-art segmentation and agglomeration such as FFN (Januszewski, Kornfeld et al. 2018) were compared against a dense connectomic reconstruction additionally including costs for synapse and type predictions and the processing of human or RoboEM skeletons (Motta, Berning et al. 2019). The RoboEM-based correction of axon split and merge errors and the attachment of spine heads only costs a fraction and therefore allows for cheap and accurate improvement of reconstructions.

Importantly, connectomic analyses differ widely in their reconstruction difficulty (Fig. 1f): for the analysis of pairs of neighboring synapses for extraction of learned synaptic configurations (Motta, Berning et al. 2019, Kornfeld, Januszewski et al. 2020), for example, axon reconstructions of about 10 μm in length are sufficient, and therefore these analyses can already now be fully automated (Motta, Berning et al. 2019, Kornfeld, Januszewski et al. 2020). Obtaining neuron-to-neuron connectomes from cortical tissue, however, requires faithful axon reconstruction for at least an additional order of magnitude of axonal length, and has so far not been possible fully automatically. Intermediate-scale connectomic analyses, aimed for example at axonal synaptic properties, demand about 50μm of error-free axonal reconstruction, that are also not yet fully automatically accessible.

Connectomic image analysis has in common that EM data are processed by CNN-based AI methods into voxel-based maps reporting plasma membranes (Jain, Murray et al. 2007, Turaga, Briggman et al. 2009, Turaga, Murray et al. 2010, Berning, Boergens et al. 2015, Lee, Zung et al. 2017, Zeng, Wu et al. 2017, Funke, Tschopp et al. 2019, Sheridan, Nguyen et al. 2021), or the similarity between pairs of image voxels (Lee, Lu et al. 2021), or the association of image voxels to the same foreground object (Meirovitch, Matveev et al. 2016, Januszewski, Kornfeld et al. 2018, Meirovitch, Mi et al. 2019). Then, segmentations are computed, and automated methods for the joining or splitting of these initial segmentation objects are currently the main focus of computational improvements (Meirovitch, Matveev et al. 2016, Lee, Zung et al. 2017, Zung, Tartavull et al. 2017, Dmitriev, Parag et al. 2018, Januszewski, Kornfeld et al. 2018, Funke, Tschopp et al. 2019, Meirovitch, Mi et al. 2019, Li, Januszewski et al. 2020, Nguyen, Jang et al. 2021). When automated approaches become insufficient, human annotation is used to solve the most challenging of these joining or splitting operations, ideally focused to those difficult locations by computational means (Plaza 2016, Haehn, Kaynig et al. 2018, Zhao, Olbris et al. 2018, Dorkenwald, Turner et al. 2019, Motta, Berning et al. 2019, Hubbard, Berg et al. 2020, Kornfeld, Januszewski et al. 2020, Scheffer, Xu et al. 2020), or by human inspection (Dorkenwald, McKellar et al. 2020, MICrONS Consortium, Bodor et al. 2021). The so far most efficient of such focused interventions (in time spent per problem solved) asks human annotators to fly along axons (Boergens, Berning et al. 2017) until another segmentation object is reached and thus the problematic location resolved (Fig. 1a, (Motta, Berning et al. 2019)).

We wondered whether the process of flight tracing along elongated (often very thin) neurites could directly be emulated in an automated fashion. For this, we used an analogy to image-based road following in autonomous driving (Pomerleau 1989, Bojarski, Del Testa et al. 2016), and developed a neural network architecture to output 3D steering signals directly from neurite-aligned 3D-EM image volumes (Fig. 1e). We (1) defined a continuous 3D steering framework; (2) defined a membrane-avoiding flight policy to recover from off-centerline positions and non-centerline-aligned orientations; (3) trained a CNN on image-steering pairs from on- and off-centerline positions and orientations in a supervised manner allowing for stable path following during recurrent inference; (4) defined a validation strategy for automated error detection (Fig. 1c-e).

In particular, we used the Bishop frame (Bishop 1975) along interpolated neurite centerline reconstructions to describe the neurites’ local directionality, defined the projection for the neurite-aligned 3D-EM input, and then used the corresponding Bishop curvatures as the target output steering signal that had to be predicted (Fig. 1d). Only during training, a neurite volume mask (obtained by segment pick-up from an oversegmentation) was used to generate off-centerline inputs and corresponding target output steering signals back towards the centerline. During inference, predicted Bishop curvatures are integrated to yield the next position and orientation. With this, we obtained an automated neurite path annotation that mimicked the process of human flight-mode annotation in 3D ((Boergens, Berning et al. 2017); Suppl. Movie 1).

Next, we investigated to what degree RoboEM could in fact replace human annotation, and which kinds of connectomic analyses would thus become fully automatable (Fig 1f). For this we started with the dense connectomic analysis of a piece of mammalian neocortex (Motta, Berning et al. 2019) in which based on automated segmentations, human annotators had been asked to resolve an automatically identified set of problem locations consuming a total of 4,000 work hours. To resolve split errors, endings of axons and spine necks had been queried, asking humans to continue if possible until the task was automatically stopped when another reconstructed object of sufficient size had been reached (Fig. 1g). To resolve merger errors, chiasmatic configurations had been detected and queried, asking humans for proper continuation from one chiasmatic exit into one of the other exits (Fig. 1g). We used RoboEM to replace these human annotations by starting RoboEM queries analogous to human annotator queries. In fact, 76% of the ending queries and 78% of the chiasma queries were automatically solved by RoboEM (Human: 74% and 94% respectively). Additionally, we used the notion that a forward continuation along a neurite should symmetrically be confirmed by tracing the same neurite location backwards for an automatic validation of RoboEM results. Restricting to RoboEM annotations in which forward and backward tracings agreed allowed avoiding most merge errors, yielding only 4% of queries with RoboEM-introduced merge errors for ending tasks and 1% for chiasma tasks, similar as for human annotations.

While this performance indicated that RoboEM could faithfully replace human annotation for a range of connectomic problems, we wanted to quantify this conclusion explicitly by using connectomic analysis itself as the metric for reconstruction success. As described above, some connectomic analyses require more reconstruction accuracy than others. In particular, we considered (1) paired same-axon same-dendrite synapse analyses aimed at measuring the learned fraction of a connectome (Motta, Berning et al. 2019); (2) spine rate analyses for identification of interneuron dendrites; (3) axonal type analysis based on the synaptic target distribution of axons. We then used three types of connectomes for comparison: (CI) the connectome obtained from the fully automated reconstruction in (Motta, Berning et al. 2019), before any focused human annotation; (CII) the connectome in (Motta, Berning et al. 2019) including 4,000 work hours of human annotation; (CIII) the connectome obtained from combining the fully automated reconstruction CI with RoboEM, yielding a fully automated and automatically proofread connectome. We then performed the three types of connectomic analyses, which we expected to increase in connectomic difficulty from requiring pairs of synapses to be faithfully linked to axons with at least 10 synapses for type definition.

When applying the analyses to the three stages of connectomes, we found that the analysis of paired synapses for quantifying potentially learned synapses in the connectome was already successful with the automated connectome state before any RoboEM-based corrections (upper bound of fraction of paired connections consistent with long term potentiation: CI 11-20%; CII 16-20%; CIII 13-19%). However, for obtaining correct spine rates for apical dendrites (CI: 0.9±0.4; CII: 1.3±0.6; CIII: 1.2±0.5 spines/μm) and the true fraction of excitatory axons defined by their spine head preference (excitatory axon fractions CI: 75%; CII: 87%; CIII: 84%) did require human or RoboEM-based corrections. Additionally, RoboEM-based corrections recovered the axonal target specificities of inhibitory axons onto apical and smooth dendrites (AD, SD), whereas in the automated state prior to RoboEM and human corrections this specificity was not detectable (One-sided Kolmogorov Smirnov test, CI: AD p=1.0%, SD p=3.6%; CII: AD p=2.7×10^−4^, SD p= 1.8×10^−3^; CIII: AD p=1.9×10^−5^, SD p=7.1×10^−4^). So in fact, when utilizing a connectomic metric of automation qualified by the difficulty of connectomic problems that can be automatically analyzed, RoboEM-based error correction shifts the automated analysis performance from simpler to more complex connectomic problems (Fig. 1f).

Next, we applied RoboEM to 3D EM data obtained using the current state-of-the-art 3D EM imaging approach for large volumes from mammalian brains (ATUM (Hayworth, Kasthuri et al. 2006) followed by multiSEM (Eberle, Mikula et al. 2015)), see (Shapson-Coe, Januszewski et al. 2021, Loomba, Straehle et al. 2022). We used a volume sized (150 μm)^3^ imaged at 4 × 4 × 35 nm^3^ resolution from a 1.3 × 1.3 × 0.25 mm^3^ sized dataset of mouse primary somatosensory cortex (Sievers, Motta, Schmidt, Helmstaedter, unpublished dataset) to quantify RoboEM performance on densely seeded axons within such a volume (Fig. 2a,b). After automated agglomeration, we seeded RoboEM at automatically detected endings of axon agglomerates, and used it to connect split axonal agglomerates (Fig. 2a). As a result, split rates of axons were reduced 7-fold (42.7 to 6.0 per mm axon path length) while only modestly increasing merger rates (3.3 to 4.5 per mm, Fig. 2b). The resulting axon reconstructions were comparable to those achieved in the local cortical SBEM dataset ((Motta, Berning et al. 2019), 5.2 split and 6.1 merge errors per mm, Fig. 2b). Similar to thin axons, thin spine necks can pose difficulties for current reconstruction pipelines. To assign synapses onto spine heads to the correct postsynaptic neuron, spine necks need to be reconstructed with high accuracy. Using RoboEM to follow automatically detected spine heads along the spine neck to the dendritic shaft of origin improved spine head attachment recall from 70% to 94%, while retaining high precision of 97% in ATUM-multiSEM data (85/91 test spine heads attached to the correct dendrite, 3/91 spine heads attached to the wrong dendrite, 3/91 spine heads unattached, of which 1 could also not be attached manually, see Supplement).

**Figure 2.**
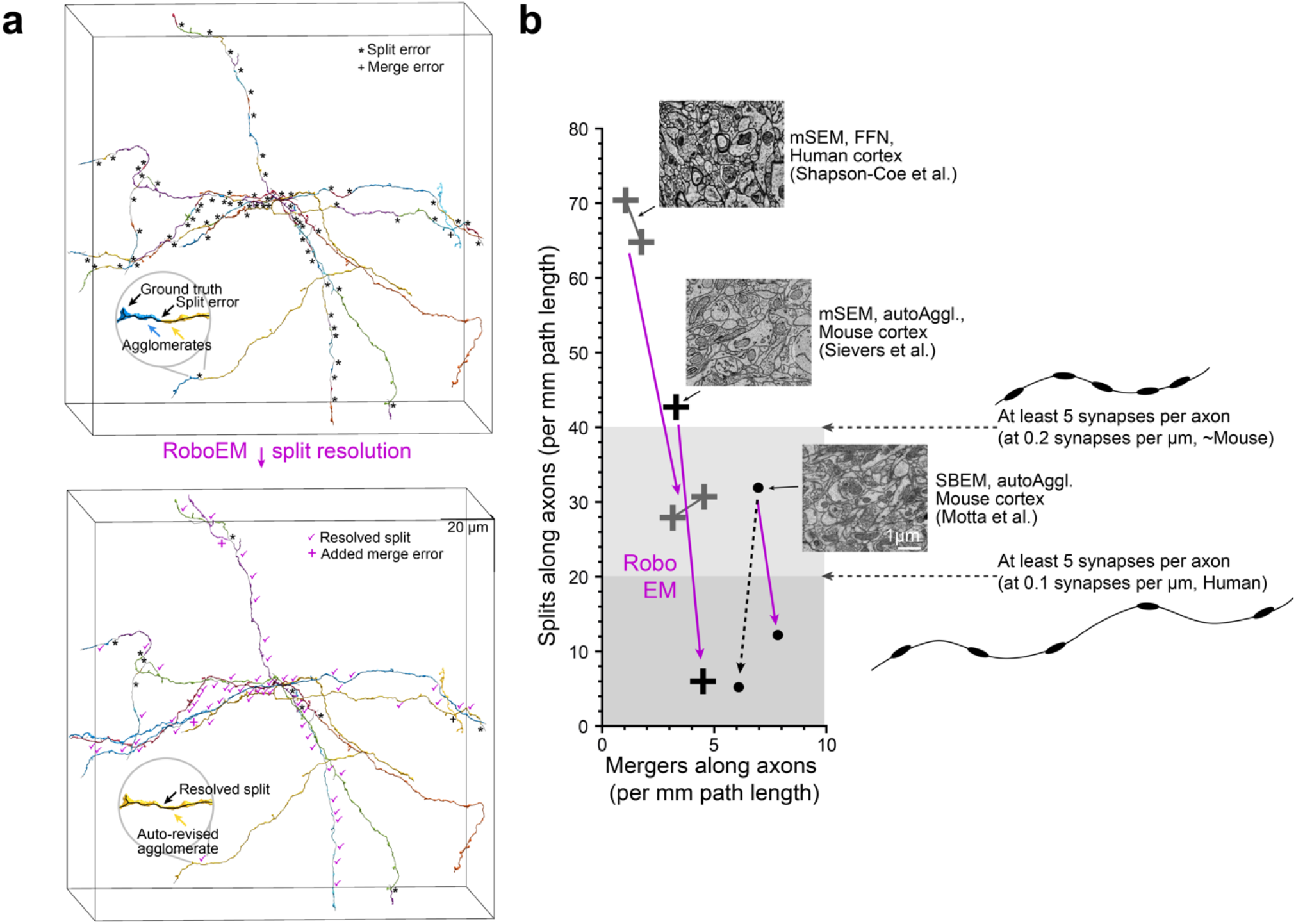
RoboEM improves state-of-the-art connectomic reconstruction results in mammalian cortex. (a) Example of dense axon sets used for unbiased calibration of reconstruction success. Note that analysis of soma-based axons highly biases towards thicker and easier parts of axons (Suppl. Fig. 3a). Agglomerates pre and post-RoboEM correction shown, left versus right. (b) Quantification of rate of axon splits and mergers, reported for randomly seeded axons. FFN-based reconstruction of human cortex (Shapson-Coe, Januszewski et al. 2021): two segmentation states analyzed (c2, c3) for which RoboEM could solve 37-40 splits per mm axonal path length (56% of all splits caused by FFN), while only adding 1.4-3.5 merge errors per mm. Note that split rate requirements depend on synapse rate along axons: for at least 5 synapses per axon, split rates below 40 per mm and 20 per mm are required for typical mouse and human axons, respectively. Moving automation into these regimes is therefore critical for most connectomic analyses (cf. Fig. 1f). RoboEM improved densely seeded axon reconstruction 7-fold for large-scale data from mouse cortex (Sievers, Motta, Schmidt, Helmstaedter, unpublished dataset). For SBEM data, RoboEM allowed the full replacement of human annotation for the employed analyses (Motta, Berning et al. 2019). The added benefit of factor 2 from manual annotations (black dashed line) was not relevant for the analyses performed in that study. Note that for fully automated analysis of local circuitry, error rates below 10 per mm are desirable.

Finally, we wondered whether RoboEM could also be applied to improve existing state-of-the-art automated segmentations which are currently being manually proofread, in particular based on flood-filling networks (FFN)-based segmentations (Januszewski, Kornfeld et al. 2018, Shapson-Coe, Januszewski et al. 2021). We evaluated RoboEM on a subvolume of size (150 μm)^3^ from (Shapson-Coe, Januszewski et al. 2021) and found that application of RoboEM on axonal endings obtained from the dense reconstructions of a flood-filling network (FFN; (Januszewski, Kornfeld et al. 2018)) segmentation solves 56% of the splits on a random set of axons traced throughout the subvolume. This reduced the split rate from 65-70 splits/mm to 28-31 splits/mm, reaching split-lengths required for automated connectomic analyses while introducing few new merge errors (merge rate increased from 1-2 mergers/mm to 3-5 mergers/mm), which are tolerable merge rates for such connectomic analyses (Fig. 2b).

These results are important for two reasons. First, evaluation of automated reconstruction performance on random axons not necessarily connected to a soma within the mm^3^-scale volume provides a representative quantification of reconstruction quality for dense connectomic reconstruction. Restriction to soma-proximal axons properly quantifies results for soma-based connectomic data, but underestimates reconstruction errors for dense axons. For example, the evaluation of FFN in human cortex, restricted to axons connected to a soma within the volume underestimates split and merge errors of random axons 4-5-fold (cf. Suppl. Fig 3a). When using soma-based reconstructions in small brain volumes(Januszewski, Kornfeld et al. 2018), error rate quantification is highly biased to the much easier-to-reconstruct dendrites and therefore cannot be interpreted for axonal reconstructions. Secondly, for the connectomic analysis of axons in cortical neuropil, it is essential to obtain an interpretable number of output synapses along axons automatically. Given a certain rate of synapses along axons, this means that there is a minimum inter-split distance that needs to be achieved in order to obtain an interpretable number of synapses per axonal segment (Fig. 2b). Importantly, the rate of synapses per axon can vary strongly between neuronal tissue types and species, such that axonal reconstructions in human cortex, for example, require at least 2-fold longer inter-split distances (i.e. 2-fold lower split error rates) than mouse cortex to achieve similar synaptic statistics per axon (Fig. 2b).

In summary, we find that our automated proofreading is able to fully replace human annotation for a number of connectomic problems, and improves currently available state-of-the-art automated segmentations. This was in particular notable for segmentations obtained using a state-of-the-art technique, flood-filling networks (FFNs (Januszewski, Kornfeld et al. 2018, Shapson-Coe, Januszewski et al. 2021)), since FFNs have a certain similarity in their design to RoboEM by recursively reconstructing individual neurites through prediction of neurite continuation from 3D EM subvolumes. RoboEM, in contrast, by focusing on centerline neurite tracing, adds a notion of growth inertia to the axonal reconstruction, which was able to solve reconstruction problems not solved by volume-based methods. Since RoboEM directly transforms EM image volumes into centerline tracings of neurites, it allows end-to-end learning of the key connectomic challenge: to follow axons over very long distances, also at locally thin stretches. Its computational cost is favorably lower than other approaches, and with this direct end-to-end strategy may allow optimization for both additional accuracy and computational efficiency, which will be the next key challenge in connectomics for exabyte-scale datasets to come (Motta, Schurr et al. 2019).

## METHODS

## See Supplementary Material

## ACKNOWLEDGEMENTS

We thank B. Staffler and M. Berning for discussions; T. Colnaghi and K. Reuter for discussions and code contributions; C. Guggenberger for compute infrastructure; V. Pinkau, N. Rzepka, J. Striebel, G. Wiese (scalable minds GmbH, Potsdam, Germany) for contribution to segmentation, agglomeration and subcellular type predictions on the multiSEM dataset.

## AUTHOR CONTRIBUTIONS

MH conceived, initiated and supervised the study; MSc conceptualized, developed and implemented all aspects of RoboEM and performed analyses; AM contributed to RoboEM development and adapted it for axon reconstruction in large scale datasets; MSi provided ATUM-multiSEM data; MSc and MH wrote the paper with contributions by all authors.

## Notes

Conflict of interest: A patent application has been filed by the Max Planck Society.

### Competing Interest Statement

A patent application has been filed by the Max Planck Society.

