## Supplementary Material for "RoboEM: automated 3D flight tracing for synaptic-resolution connectomics"

### MATERIALS AND METHODS

#### EM image datasets, segmentations, RoboEM training, validation and test sets

RoboEM was developed and tested on a 92.6 x 61.8 x 94.8 $\mu m^3$ SBEM dataset (Denk & Horstmann, 2004) from layer 4 primary somatosensory cortex of a 28 days old mouse previously densely reconstructed by (Motta et al., 2019). The tissue was conventionally en-bloc stained (Briggman, Helmstaedter, & Denk, 2011) and imaged at 11.24 x 11.24 $nm^2$ and nominal cutting thickness of 28 nm. Training and validation axons were sampled from a set of axons seeded by means of presynaptically classified segments obtained using SynEM (Staffler et al., 2017) and skeleton traced by student annotators. To also acquire a volume reconstruction of training axons, segments of the oversegmentation obtained using SegEM with parameters as set for the whole-cell segmentation of the cortex dataset (Berning, Boergens, & Helmstaedter, 2015) were picked up and combined. The volume mask was also used to iteratively optimize the interpolated skeleton tracing to yield a better centerline approximation. A set of 14 axons with 1.2 mm path length was used for training, and up to 700,000 weight updates corresponding to ~260 epochs were run. The validation set consisted of 13 axons with 1.4 mm path length, where branches with less than 5 µm were excluded for better heuristic error detection. Results on the validation set over the course of training are depicted in Suppl. Fig 1b. A third set of 10 axons with 1.7 mm path length seeded from a $(2.5\ \mu m)^3$ bounding box was used as a test set, cf. Fig. 2b, Suppl. 1c. These are the same axons as previously used to evaluate human and semi-automated segmentations (Boergens et al., 2017; Motta et al., 2019). For automated spine head attachment on this dataset the axon-trained RoboEM was evaluated on a random subset of 50 spine heads previously attached by human annotators, as well as on the set of spine heads previously used as test set in (Motta et al., 2019).

Next, RoboEM was trained, tested and applied on a subset of a 1.3 x 1.3 x 0.25 $mm^3$ dataset from the barrel cortex of a 28 days old mouse stained following the protocol by Hua et al. (Hua, Laserstein, & Helmstaedter, 2015) with small modifications, sectioned at 35 nm using ATUM (Hayworth, Kasthuri, Schalek, & Lichtman, 2006) and imaged at 4 x 4 $nm^2$ with a multibeam scanning electron microscope (Eberle et al., 2015) (Sievers, Motta, Schmidt, Helmstaedter, unpublished dataset). The segmentation and agglomeration applied to the dataset at a resolution of 8 x 8 x 35 $nm^3$ (downsampled in x-y by a factor of 2) were developed in collaboration with scalable minds GmbH partly based on published approaches. In brief, a 3D U-Net (Ronneberger, Fischer, & Brox, 2015) was used to predict per cardinal axis voxel affinities following (Lee, Zung, Li, Jain, & Seung, 2017) from which a watershed-based oversegmentation was generated. The segments from the oversegmentation were then combined using hierarchical agglomeration as proposed in (Funke et al., 2019). In addition, neurite type predictions, blood vessel and nuclei detection were incorporated into the agglomeration to further reduce merge errors. Neurite type predictions were also used for spine head detection, which was the basis for RoboEM-based spine head attachment. For RoboEM training on axons, a training set was acquired based on a set of 10 soma seeded axons from Layer 4, for which student annotators picked up segments from the oversegmentation to acquire a volume reconstruction. Here, we then used Kimimaro (Silversmith, Bae, Li, & Wilson, 2021) followed by subsampling and B-spline interpolation to extract centerline skeletons from the volume reconstruction. This yielded a training set with a total of 21 mm axon path length. As axon validation set a random 10 out of 20 axons from a Layer 4 seeded bounding box of size (2 μm)$^3$ were traced within a (50 μm)$^3$ bounding box yielding 1 mm path length. For the axon test set, another (1.5 μm)$^3$ bounding box within layer 4 was densely annotated and a random subset of 5 axons were traced within a (150 μm)$^3$ bounding box yielding 1.7 mm path length. A separate RoboEM model was trained on spine attachment. For this training and validation set were generated from a set of 20 (5 μm)$^3$ bounding boxes sampled within layer 4 and annotated for spine heads. A subset of around 1000 spine heads were volume annotated at 4 x 4 x 35 nm$^3$ by student annotators from the spine head through the spine neck up to the dendritic trunk. Here, we again used Kimimaro (Silversmith et al., 2021) to extract centerline skeletons from the volumetric masks for training yielding 2 mm of spine neck tracings. Additionally, a random subset of 76 spine heads with 0.2 mm path length from the 20 bounding boxes was skeleton traced to serve as validation set. The evaluation was done on another randomly selected (5 μm)$^3$ bounding box containing 91 densely annotated spine head to dendritic trunk skeleton tracings.

For the evaluation of RoboEM-based error correction of another state-of-the-art segmentation, we applied RoboEM to a recently published mm$^3$-scale multiSEM dataset with voxel size 4 x 4 x 33 nm$^3$ (Shapson-Coe et al., 2021) segmented and agglomerated using flood-filling networks (FFN; (Januszewski et al., 2018)). Specifically, we focused on a (150 μm)$^3$ bounding box containing 6.5 mm of axon path length of the provided ground truth skeleton tracings. The published ground truth skeleton tracings in this box were then used to evaluate FFN on soma-seeded axons, cf. Suppl. Fig 3a. To finetune RoboEM on this dataset, we generated ground truth skeleton tracings of dense seeded axons by sampling a bounding box of size (2.5 μm)$^3$ within a centered (15 μm)$^3$ bounding box, annotating all processes in this bounding box and then sampling a random subset of 5 axons, which were traced throughout the (150 μm)$^3$ bounding box yielding 1.25 mm path length. Here, skeleton annotations were already done with high precision along the centerline such that no postprocessing was necessary. The volumetric neurite mask needed for training was generated by means of segment pick-up from the c3 FFN segmentation, cf. (Shapson-Coe et al., 2021). The best RoboEM model checkpoint, based on the validation set from the axon training in the other multiSEM dataset (Sievers et al., unpublished dataset), was then used as initialization and RoboEM was trained for another ~2 million gradient updates until converging in terms of reset-based error rates on the training axons. For the evaluation of FFN and RoboEM another bounding box of size (1.5 μm)$^3$ was annotated for a random subset of 5 axons yielding 1.4 mm path length.

#### Neurite flight reconstruction

We phrase the problem of neurite reconstruction from a 3D-EM volume as a centerline reconstruction task, in which a neurite is represented by a sequence of visited points. A convolutional neural network (CNN; (LeCun et al., 1989)) is trained on the task of predicting the local neurite continuation from a neurite-centered and –aligned 3D-EM sub-volume, similar to human annotation flight mode (Boergens et al., 2017), cf. Suppl. Fig 1. Integration of the predicted neurite continuation yields a new position and orientation, which is used to generate the subsequent CNN input. Iterative application of this procedure turns a starting location and orientation into a neurite skeleton reconstruction using only 3D-EM data and without intermediary steps, such as volume segmentation.

The input to the CNN consists of a 96 x 96 x 16 Vx neurite-centered and –aligned 3D-EM sub-volume that covers, e.g. for axons, a field of view of $\sim 1 \times 1 \times 0.7\ \mu\mathrm{m}^3$ (cf.

Suppl. Fig. 1a). The third (Z) dimension corresponds to the current flight direction, while the current position is made the center of the 4th Z-plane. The field of view is thus asymmetric along the Z direction with more contextual information available in forward than in reverse flight direction. Empirically, this allows for better steering towards the 'exit' within axonal varicosities. The EM data is projected onto neurite-aligned planes by means of trilinear interpolation.

For the CNN architecture we use 7 3D strided convolutional layers, followed by a dropout layer (dropout rate: 0.5), 3 fully connected layers and a last linear layer that estimates the two steering commands and the distance to membrane, cf. Suppl. Fig 1a. The two steering commands are the Bishop curvatures described in more detail below. The 2D CNN architecture used for image-based road following in (Bojarski et al., 2016) served as a starting point for the proposed architecture in this work. As nonlinearity we tested rectified linear units (ReLU; (Glorot, Bordes, & Bengio, 2011)) and exponential linear units (ELU; (Clevert, Unterthiner, & Hochreiter, 2015)) and found ELU to work better.

For the mathematical formulation of the network's input and output the Bishop frame and the associated Bishop curvatures are particularly suitable (Bishop, 1975) – cf. Fig. 1d and Suppl. Fig. 1a. The Bishop frame is a local orthonormal coordinate system spanned by a tangential vector and two normal vectors and obeys a rotation minimizing property disallowing any twists around the tangential vector. In this work, we define the neurite aligned projection planes for the network's input by means of the Bishop normal vectors, while the network's flight direction corresponds to the tangential vector. The evolution of the Bishop frame unit vectors is coupled via the Bishop curvatures, one for each normal vector direction. In this work, we use the Bishop curvatures as the network's output for steering. Each of the two Bishop curvatures resembles a signed curvature for the corresponding normal vector direction. The signed curvature has also been used as steering command for image-based road following, albeit only for one steering direction corresponding to planar curves (Bojarski et al., 2016). The task of predicting the neurite continuation in the form of Bishop curvatures can be interpreted as fitting a parabola to the neurite's centerline. The curvature vector, i.e. the sum of Bishop curvatures multiplied by their corresponding Bishop normal vector, determines direction and magnitude of bending of this parabola, cf Fig 1d and Suppl. Fig 1a.

#### B-Spline Interpolation and RoboEM step size

To get a continuous representation of neurite branches from the sparsely placed nodes from human skeleton reconstructions, we use degree 4 B-spline interpolation (Piegl & Tiller, 1996) yielding a curve $\vec{\gamma} \in C^3$. We reparametrize the curve to get a curvature adaptive step size $\|\dot{\vec{\gamma}}\|\,\Delta t$ of

$$\|\dot{\vec{\gamma}}\|\,\Delta t = f \cdot \frac{d}{1 + \frac{p}{2}\,\kappa},$$

where $\kappa$ is the curvature, and $p = 1\ \mu\mathrm{m}$ the physical size of the projection plane and $f$ is a step size factor set to $f = 1$ for training and can be adjusted to higher values for inference. While for most evaluations in this work, we also used $f = 1$ during inference, step sizes can be increased up to $f = 5$ without risking worse performance of RoboEM, cf. Suppl. Fig 1c. A default step size $d$ matching the smallest dimension of a voxel, such as $d = 11.24\ \mathrm{nm}$ in case of the L4 SBEM dataset (Motta et al., 2019) and with $f = 1$ ensures that no voxels up to a radius of half the projection plane size in the projection plane at the current position of the CNN are skipped. To keep the notation uncluttered subsequent formulas are expressed in terms of the arc length parametrized curve $\vec{\gamma}(s)$ indicated by the parameter $s$.

#### Bishop frame

The Bishop frame (Bishop, 1975) consists of three orthonormal vectors, namely the tangential vector $\vec{t}$, in this work also termed flight direction, and two normal vectors $\vec{n}_1, \vec{n}_2$. The evolution equations of the Bishop frame and the curve $\vec{\gamma}$ read as follows:

$$\frac{\mathrm{d}}{\mathrm{d}s}\vec{\gamma} = \vec{t}$$

$$\frac{\mathrm{d}}{\mathrm{d}s}\vec{t} = k_1\vec{n}_1 + k_2\vec{n}_2 \equiv \vec{k}$$

$$\frac{\mathrm{d}}{\mathrm{d}s}\vec{n}_1 = -k_1\vec{t}$$

$$\frac{\mathrm{d}}{\mathrm{d}s}\vec{n}_2 = -k_2\vec{t}\,.$$

Here, $k_1$ and $k_2$ are the Bishop curvatures associated to the normal vector $\vec{n}_1$ and $\vec{n}_2$ respectively, and we defined the curvature vector $\vec{k}$ (cf. Fig. 1d, Suppl. Fig 1a), which obeys $\|\vec{k}\| = \sqrt{k_1^2 + k_2^2} = \kappa$, where $\kappa$ is the curve's curvature. Note that flips and rotations around the tangential vector applied to both normal vectors and Bishop curvatures are invariance transformations that leave $\vec{k}$ and thereby the evolution of $\vec{\gamma}$ and $\vec{t}$ unchanged. Application of these transformations to the normal vectors defining the CNN's input, and to the Bishop curvatures, i.e. the CNN's output, defines an equivariance, on which we train the CNN by data augmentation. Additionally, we can also flip the flight direction, which corresponds to a time reversal symmetry. Note that on the neurite centerline this time reversal and tangential vector flip does not affect the Bishop curvatures, while for the flight policy derived in the next section this symmetry is broken. Note that the rotation minimizing Bishop frame has weaker requirements for $\vec{\gamma}$ than the Frenet-Serret frame often used as local coordinate system in differential geometry of parameterized curves. Namely, the Bishop frame of a parametrized curve in 3D Euclidean space does not require $\kappa \neq 0$, and only requires $\vec{\gamma} \in C^2$ instead of $\vec{\gamma} \in C^3$ (Bishop, 1975). Additionally, initial conditions in form of start position and Bishop frame orientation together with the Bishop curvatures uniquely define the centerline curve $\vec{\gamma}$ (Bishop, 1975), which makes the Bishop curvatures particularly suitable as steering command, as it allows to reconstruct the curve by means of above evolution equations. The integration of above Bishop equations during inference was performed with either of two methods: 1) forward Euler method followed by Gram-Schmidt orthonormalization to maintain an orthonormal basis for the Bishop frame, or 2) analytically derived evolution equations for a Bishop frame along a parabola. Empirically, we found the latter to work better especially for larger step size factors $f$, cf. Suppl. Fig. 1c.

#### Flight policy

To derive a flight policy, we start with the Taylor series expansion up to second order $\vec{\gamma}_T$ for the known neurite centerline curve $\vec{\gamma}(s)$ with corresponding Bishop frame $\vec{t}, \vec{n}_1, \vec{n}_2$ and curvature vector $\vec{k}$, and the Taylor series expansion up to second order $\underline{\vec{\gamma}}_T$ of the unknown off-centerline, off direction curve $\underline{\vec{\gamma}}(s)$, with $\underline{\vec{t}}, \underline{\vec{n}}_1, \underline{\vec{n}}_2, \underline{\vec{k}}$:

$$\vec{\gamma}_T(s) = \vec{\gamma} + s\,\vec{t} + \frac{s^2}{2}\vec{k}$$

$$\underline{\vec{\gamma}}_T(s) = \underline{\vec{\gamma}} + s\,\underline{\vec{t}} + \frac{s^2}{2}\underline{\vec{k}}\,.$$

Note that the second order Taylor series (parabola) approximation only depends on the current position $\vec{\gamma}$, the flight direction $\vec{t}$ and the curvature vector $\vec{k}$. Hence, the prediction of $\vec{k}$ can be interpreted as fitting a parabola to the neurite of interest.

Knowing the current off position $\underline{\vec{\gamma}}$ and off direction $\underline{\vec{t}}$, as well as the current closest position on the neurite centerline $\vec{\gamma}$ with correct neurite aligned orientation $\vec{t}$ and local shape of the curve in terms of $\vec{k}$, i.e. correct steering for the on centerline case, we would like to find suitable corrected steering $\underline{\vec{k}}$, which converges back to the centerline within some distance $s_c$, i.e. which minimizes the future distance $\|\vec{\Gamma}_T\|$ defined by:

$$\underbrace{\vec{\gamma}_T - \underline{\vec{\gamma}_T}}_{\vec{\Gamma}_T} = \underbrace{\vec{\gamma} - \underline{\vec{\gamma}}}_{\vec{\Gamma}} + s\,\underbrace{(\vec{t} - \underline{\vec{t}})}_{\vec{\mathrm{T}}} + \frac{s^2}{2}\underbrace{(\vec{k} - \underline{\vec{k}})}_{\vec{\mathrm{K}}}\,.$$

We therefore require:

$$\nabla_{\underline{\vec{k}}}\left\|\vec{\Gamma}_T(s_c)\right\|^2 = 0\,,$$

where $\nabla_{\underline{\vec{k}}} = \underline{\vec{n}}_1\partial_{\underline{k}_1} + \underline{\vec{n}}_2\partial_{\underline{k}_2}$. Defining the projection operator $\mathcal{P}_{\underline{\vec{n}}_1,\underline{\vec{n}}_2} = \underline{\vec{n}}_1 \otimes \underline{\vec{n}}_1 + \underline{\vec{n}}_2 \otimes \underline{\vec{n}}_2$, where $\otimes$ denotes an outer product, the solution for $\underline{\vec{k}}$ from above equation reads as follows:

$$\underline{\vec{k}} = \mathcal{P}_{\underline{\vec{n}}_1,\underline{\vec{n}}_2}\left(\frac{2}{s_c^2}\left[\vec{\Gamma} + s_c\vec{t}\right] + \vec{k}\right).$$

Here, $s_c$ is a free parameter of this flight policy. Trajectories for different values of $s_c$ are plotted in Suppl. Fig. 1b. For this work, during training, we set $s_c$ to the current distance to membrane along the flight direction and thereby train RoboEM on a membrane avoidance strategy.

#### Direction prediction with Monte-Carlo Dropout

For the direction prediction, as e.g. needed for spine head attachment where only an initial position is given, we sample for instance $I = 256$ roughly equidistant orientations $\vec{t}_i$ and run Monte-Carlo Dropout with e.g. $M = 128$ samples. From the sampled Bishop curvature predictions $\vec{k}_{im}$, we compute the mean curvatures $\langle\kappa\rangle_i$ and the covariances of the Bishop curvatures $cov_i$ per orientation $i$ over the Monte-Carlo samples. The uncertainty estimate $u_i$ is then taken as the square-root of the largest eigenvalue of the covariance matrix divided by the mean curvature

$$u_i = \sqrt{max(eigvals(cov_i))} / \langle\kappa\rangle_i \ .$$

Using cosine similarity for neighboring angles within 30 degrees, we construct a weighted average of uncertainties $\hat{u}_i$ as

$$\hat{u}_i = \frac{u_i}{2} + \frac{1}{2}\sum_j \widehat{w}_{ij}\, u_i$$

with weights $\widehat{w}_{ij}$

$$\widehat{w}_{ij} = {w_{ij}} \Big/ {\textstyle\sum_j w_{ij}}$$

$$w_{ij} = \begin{cases} \langle \vec{t}_i | \vec{t}_j \rangle^{16} & \text{if } \arccos\left(\langle \vec{t}_i | \vec{t}_j \rangle\right) < 30° \\ 0 & \text{otherwise} \end{cases}$$

to give more stable predictions. The orientation with minimal averaged uncertainty $\hat{u}_i$ is then the first orientation candidate. For spine head attachment in the multiSEM dataset (Sievers et al., unpublished dataset), we use a second candidate that is >110 degrees from the first and has again minimal averaged uncertainty among the remaining orientations.

#### Training and inference

For training above described CNN architecture on the neurite following task, we implement the CNN in TensorFlow (Abadi et al., 2016) and train with a mini-batch size of 128 using RMSProp (Tieleman & Hinton, 2015) with momentum (Rumelhart, Hinton, & Williams, 1986; Sutskever, Martens, Dahl, & Hinton, 2013) or Adam (Kingma & Ba, 2015) on minimizing the mean squared error plus a L2 regularization loss term on the weights. Weights for layers with ReLU or ELU activations were initialized following (He, Zhang, Ren, & Sun, 2015), while the last layer was initialized following (Glorot & Bengio, 2010).

Similarly to the finding of (Pomerleau, 1989) for image-based road following, we also find that training on the neurite centerline alone does not yield good generalization performance during inference. Note that during inference current position and orientation depend on past network decisions. Therefore, even small errors in past network's steering predictions result in off-centerline positions. To achieve stable path following, we train the CNN on off-centerline positions and off directions with correspondingly adjusted steering leading back to the neurite centerline. The mapping from a particular off position, off direction state to the adapted steering is referred to as flight policy.

For this work, we derived a greedy flight policy only based on local information available to the CNN within the finite field of view. The distance within which to converge back to the centerline is a free parameter of this flight policy. Suppl. Fig. 1b shows trajectories obtained using different convergence distances for off-center positions within a synaptic bouton. Evidently, when starting closer to the plasma membrane, more aggressive steering with shorter convergence distance is required to stay within the neurite, while for larger distances to the plasma membrane a gentler steering is affordable and better preserves alignment to the centerline. Hence, we use a dynamic convergence distance that is set to the distance to the plasma membrane along the flight direction. This induces the notion of obstacle, i.e. membrane, avoidance and empirically performs better than a constant value for the convergence distance. Additionally, we used the distance to plasma membrane as an auxiliary loss term during training.

To reconstruct neurites during inference, the steering predictions of the CNN for a position and orientation are integrated to a new position and orientation used to generate the subsequent input as sketched in Suppl. Fig. 1a (normal inference). For the random rotation inference mode, cf. Suppl. Fig. 1a, we additionally perform a random rotation around the tangential after the integration step, which decorrelates consecutive inputs and comes at neglectable computational cost. In both inference modes, given a start position and orientation the recurrent application of the CNN yields a trace of visited points corresponding to the network's prediction of the neurite centerline.

While, for an error correction framework like FocusEM (Motta et al., 2019), both start position and orientation can be provided, for some use cases the orientation might not be available. Specifically, for the task of attaching spine heads to their corresponding dendrites, it can be less obvious how to compute the start orientation. To apply RoboEM on spine neck tracing tasks given only the start positions, we run stochastic forward passes (Monte-Carlo Dropout) through the dropout layer yielding a sampled distribution of predictions. Following (Gal & Ghahramani, 2016), we can estimate RoboEM's uncertainty from these samples. The uncertainty can then be used to select a start orientation from a list of candidate orientations. As there is only one dropout layer after the convolutional layers in our architecture of RoboEM, the computation for different dropout masks is shared up to this point and the overhead of Monte-Carlo (MC) Dropout is negligible (<2% in terms of FLOPS for 128 MC samples). Additionally, estimating uncertainties can also be convenient for other use cases of RoboEM.

#### Model selection from validation set

To choose a model from a pool of trained models with different learning rates, field of views, etc. we evaluated RoboEM on a validation set consisting of human skeleton annotations without any segmentation. Specifically, RoboEM was evaluated on linear neurite branches by running recurrent inference starting from both sides. In case RoboEM reaches a first set of thresholds (see Suppl. Table 1) concerning the distance or the angle w.r.t. the ground truth tracing, we consider the tracing to be 'experimental', i.e. the progress along the ground truth tracing is temporarily not considered until the first set of thresholds is not exceeded anymore. If a second set of thresholds (see Suppl. Table 1) is exceeded the tracing is classified as erroneous and a reset to the closest point on the ground truth before the tracing turned 'experimental' is performed. Tracing is stopped, when the closest point on the ground truth is at the other end of the axonal branch and the tracing is at that time classified as correct.

| **Tracing considered** | **Distance $\Gamma$ to ground truth** | **Angle $\theta$ to ground truth** |
|---|---|---|
| **correct** | $\Gamma \leq$ 360 nm | $\theta \leq 90°$ |
| **experimental**<br>→ progress along ground truth not considered | 360 nm < $\Gamma \leq$ 800 nm | $90° < \theta \leq 160°$ |
| **wrong**<br>→ reset to ground truth | $\Gamma$ > 800 nm | $\theta > 160°$ |

**Suppl. Table 1**

To relate resets caused by steering errors to merge and split error rates, we consider each reset both a merge error into a wrong process and a split error due to not continuing the neurite of interest. Hence, we count each reset as two errors and report this as a reset-based error rate. While this cannot be used to directly compare with segmentation error rates, it serves as a metric for model selection.

For model selection on the SBEM L4 dataset (Motta et al., 2019), we find that averaged over all models and training iterations the random inference mode outperforms its normal mode counterpart by 33% (range: 12-52%). The best performing model on the validation set uses only EM data as input, has an ELU activation function, was trained for 700,000 training iterations and yields, based on heuristic error detection via resets, 16.4 errors/mm. The manual inspection of RoboEM tracings in the raw data yields 1 false positive and 0 false negative resets (precision 96%, recall 100%). The true error rate on this validation set is 15.7 errors/mm.

We also test on a random set of 10 axons with 1.7 mm path length previously used for the quantification of the human error rate (Boergens et al., 2017). On this test set RoboEM yields 58.1 errors/mm via the heuristic error detection. Manual inspection reveals 4 false positive and 2 false negative resets (precision 96%, recall 98%) leading to a true error rate of 55.6 errors/mm.

##### RoboEM as error correction framework

To show that RoboEM enables fully-automated connectomic reconstructions of relevant volumes when used as an error correction framework, we reran the dense reconstruction of (Motta et al., 2019) using RoboEM as a direct replacement for human annotations within the FocusEM framework. Specifically, we trained RoboEM on axons as previously described and applied it to axon and spine neck reconstruction.

We then ran RoboEM on all detected spine heads that previously required human annotations (38% of all detections consuming around 900 working hours of annotators) and found that this allows to improve recall from automated methods as used in (Motta et al., 2019) from 58% to 70%, while precision decreased from 93% to 85%, which compares to 96%/91% when using human annotations. In detail, we used the start positions from the spine head detection also provided to human annotators and first ran the direction prediction via Monte Carlo Dropout as previously described. We then applied the recurrent inference mode of RoboEM for the first candidate direction with lowest uncertainty and traced until a dendritic trunk defined by a dendrite mask was reached or a path length threshold was exceeded. Notably RoboEM has not been retrained on spine necks for this dataset, such that further improvements can likely be achieved by assembling a dedicated training set for spine necks as has been done for the multiSEM dataset.

For the correction of split and merge errors in the axon reconstruction, which previously consumed 3000 working hours of annotators, we supplied start positions and directions from the FocusEM framework to RoboEM. For split error resolution, RoboEM was iteratively run on stretches of 1.5 µm and each stretch was validated by running RoboEM backwards. Only if the validation was successful, the tracing continued with the next stretch until a known axon agglomerate or the end of the dataset was reached. The resulting skeleton tracings could then be used analogous to human annotations. For merge error resolution, the stop criterion was based on a bounding box around the merge error – again analogous to human – and tracings were accepted if validation in backward direction yielded the same skeleton reconstruction with some error tolerance. Split error resolution was run once (~128,000 ending queries) and flight paths, for which a new agglomerate was found and the full path length was validated were incorporated (~60% of all ending queries). Next, three rounds of merge error resolution were run until less than 50% of chiasmata were solved by RoboEM annotations (first round: ~8,100 chiasmata, ~4,400 solved; second round: 4,100 chiasmata, 3,300 solved; third round: ~1,000 chiasmata, ~300 solved). Finally, partially validated flight paths from the first split error resolution were added to the agglomerates and segments that only overlap with flight paths from a single agglomerate were added to the respective agglomerates. The final axon reconstruction for this fully automated approach yielded 12 split errors/mm and 8 merge errors/mm on the test set axons comparing to 5 split errors/mm and 6 merge errors/mm for the semi-automated reconstruction, cf. Fig 2b.

Using the fully automated reconstruction acquired by previous automated methods and RoboEM, we reran parts of the biological analysis as done by (Motta et al., 2019). Most important figures for the three reconstruction states, such as number of axons, synapses etc. are summarized in Suppl. Table 2. Results of the paired same-axons same-dendrite analysis are shown in Suppl. Fig. 2a. While there are many more synapse pairs recovered for the reconstruction states obtained by human and RoboEM-based correction (Number of same axon same dendrite pairs (I) n = 993, (II) n = 5290, (III) n = 3982), the resulting fractions of paired connections consistent with long term potentiation (LTP) is similar across states ((I) 11-20%, (II) 16-20%, (III) 13-19%). In contrast, the spine densities prior to corrections are 1.4-fold lower for apical dendrites than after human proofreading, while RoboEM-based correction yields 1.08-fold lower spines per µm, cf. Suppl. Fig 2b. Further, when distinguishing excitatory and inhibitory axons with ≥10 synapses based on their fraction of primary spine innervation, in the automated agglomeration there are 62% fewer axons reaching the synapse number threshold compared to the final reconstruction using human annotations. Additionally, the resulting proportion of excitatory versus inhibitory axons is skewed yielding only 75% excitatory axons (estimate from (Motta et al., 2019): 87%;

cf. Suppl. Fig 2c). RoboEM-based correction of split and merge errors not only recovers 97% of axons reaching the synapse number threshold, but also yields a better estimate of the proportion of excitatory to inhibitory axons of 84%, cf. Suppl. Fig 2c. Finally, target specificities of axons where evaluated across reconstruction states against a binomial null model. While the overall fractions of specific axons are similar across reconstruction states (fraction of exc. axons specific for false detection rate thresholds 5-30%: (I) 9-35%, (II) 9-33%, (III) 6%-30%; fraction of inh. axons specific: (I) 29-55%, (II) 38-62%, (III) 45-69%, cf. Suppl Fig 2d), the automated reconstruction prior to human or RoboEM-based correction fails to detect apical and smooth dendrites specificities of inhibitory axons, cf. Suppl. Fig 2d.

| | **(I) autoAggl.** | **(II) autoAggl. + Human corrections** | **(III) autoAggl. + RoboEM corrections** |
|---|---|---|---|
| # axons ≥10 synapses | 2623 | 6979 | 6795 |
| # excitatory axons ≥10 synapses | 1904 | 5894 | 5599 |
| # inhibitory axons ≥10 synapses | 651 | 893 | 1058 |
| # synapses onto soma | 4668 | 4742 | 5101 |
| # synapses onto whole cells | 33164 | 45706 | 43923 |
| # synapses onto apical dendrites | 9816 | 14090 | 12985 |
| # synapses onto smooth dendrite | 15397 | 17908 | 18554 |
| # synapses onto AIS | 547 | 615 | 689 |
| # synapses onto other | 96230 | 149431 | 138910 |

**Suppl. Table 2**

#### Split and merge error evaluation

For the evaluation of split and merge errors of agglomerations before and after RoboEM-based error correction (cf. Fig 2 and Suppl. Fig 3a), we detected merge errors which extended further than 2.2 µm from the ground truth and manually verified that this heuristic accurately detects merge errors. For agglomerations prior to

RoboEM corrections, each merge error was counted as 1/2 and divided by the ground truth path length to yield the merge error rate. This is because each merge error usually connects two neurites and counting them as 1 error per neurite instead of 1/2 would overestimate the total amount of merge errors. For sparse evaluations of RoboEM-based error corrections limited to agglomerates that overlap with the ground truth, as done in both multiSEM datasets evaluated in this work, additional mergers introduced by RoboEM were counted as 1 instead of 0.5, which accounts for mergers from agglomerates not overlapping with ground truth and therefore not observable in a sparse evaluation. For split errors, we restricted the set of agglomerates to be evaluated to those agglomerates that overlap more than 2.5 μm with the ground truth. We introduced this overlap threshold in order to avoid domination of split errors by many small agglomerates or unagglomerated segments along thin stretches of axons. Note that despite this overlap length threshold around 90% of the ground truth was still covered.

For the multiSEM datasets RoboEM-based correction was restricted to ending resolution, for which endings were extracted from skeleton representations of those agglomerates that overlap with the ground truth annotations. In addition to the RoboEM validation strategy, we also made use of subcellular type predictions to decide whether a RoboEM tracing should connect two agglomerates. Note that type predictions were also used by FFN to avoid merge errors across subcellular types (Shapson-Coe et al., 2021). After agglomerates have been reconnected using RoboEM tracings, the resulting agglomeration state was evaluated as before.

##### Computational cost

To estimate computational costs, we benchmarked RoboEM on the ~128,000 ending tasks from the first set of ending detections in (Motta et al., 2019). The step size factor was set to $f = 5$ and analytically derived equations for the Bishop frame along a parabola were used to integrate steering predictions. This is in contrast to other evaluations presented in this work, which used $f = 1$ and integration using the forward Euler method (for the validity of increasing step size and hence throughput, cf. Suppl. Fig. 1c). The total runtime on a single node using 32 cores (Intel(R) Xeon(R) Gold 6130 CPU, 2 sockets), a single Tesla V100 GPU (PCIE-16GB) and less than 128GB RAM was 13.6 hours for the reconstruction of around 2.1 meters of axons (including backward validation tracings, a total of 64 million CNN inferences), and hence a reconstruction speed of around 160 mm/h ( ~1300 steps/s, average step size of 33 nm). From this and previous RoboEM runs with $f = 1$ and integration using the forward Euler method, we extrapolated to yield a total runtime of 27.9 hours for axon and spine neck tracing (4.3 meters including path length for validations), and 7.6 hours for initial direction prediction (on ~138,000 spine heads) on a single node for the automated error correction of the reconstruction state from (Motta et al., 2019) before human interventions yielding a total of 35 single GPU node hours. For flood-filling networks the 6.964 GPU node hours were multiplied by the 1000 (NVIDIA P100) GPUs (Januszewski et al., 2018). For (Motta et al., 2019), the dense reconstruction including segmentation, agglomeration, type and synapse prediction and processing of human skeleton reconstructions took 101 hours on 24 nodes with 16 CPU cores each. At the time of writing, Amazon EC2 (x2gd.8xlarge instance) costs per CPU core are at 0.0835 USD/h and a single T4 GPU node with 32 cores and 128GB RAM (g4dn.8xlarge) costs 2.176 USD/h. Since T4 GPUs only have 8 TFLOP (single precision) in comparison to 14 TFLOP on V100 GPUs and 11 TFLOP on P100 GPUs, we adapted the costs accordingly. For all methods the costs were multiplied by the ratio of the respective dataset sizes and normalized to the 2.7 meters of neurites (Motta et al., 2019), cf. Suppl. Table 3, resulting in the compute cost estimates in Fig. 1h, i. Note that at the time of writing 1 USD ≈ 1 EUR.

| **Approach** | **Hardware** | **Runtime** | **CPU core hours** | **GPU node hours** | **Dataset size factor** | **Cost** [EUR/m] |
|---|---|---|---|---|---|---|
| **FFN** (Januszewski et al., 2018) | 1000x NVIDIA Tesla P100 nodes *(2.992 USD/h/P100)* | 6.964 h | - | 6964 | 0.5 | 3904 |
| **Dense reconstruction** (Motta et al., 2019) | 24x CPU nodes with 16 CPU cores each @ 16 GB RAM / core *(0.0835 USD/h/core)* | 101 h | 38.8 k | | 1 | 1200 |
| **RoboEM** | 1x NVIDIA Tesla V100 node with 32 CPU cores @ 128GB RAM *(3.808 USD/h/V100)* | 35 h | - | 35 | 1 | 49 |

447 **Suppl. Table 3**

### FIGURE LEGENDS

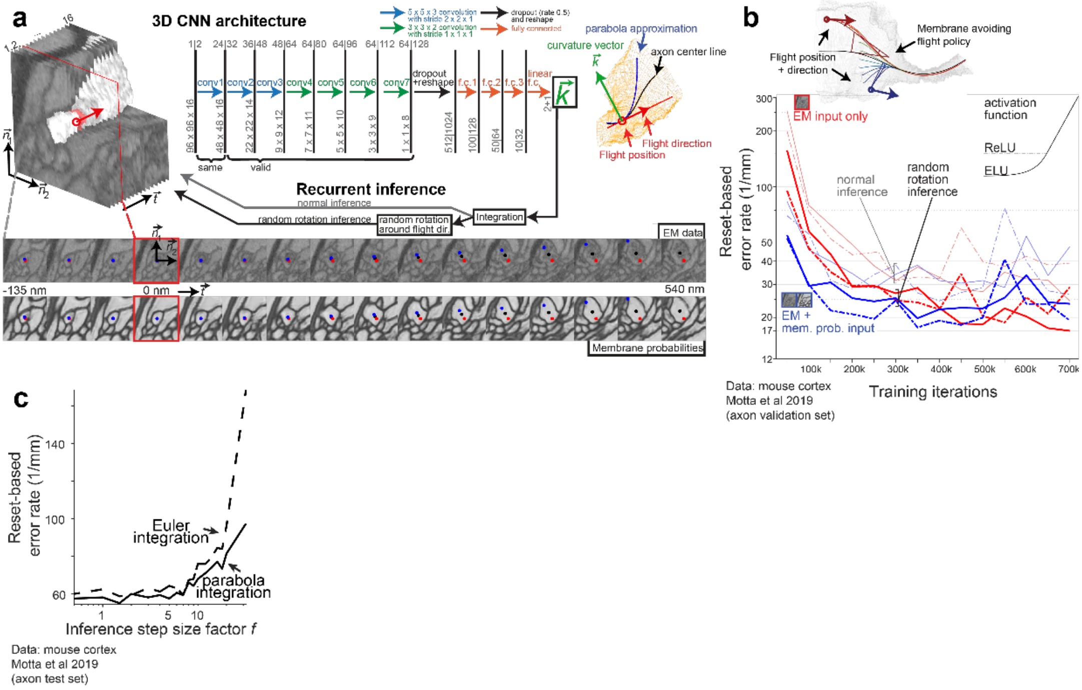


**Suppl. Figure 1. Automated flight reconstruction.** (a) Exemplary 3D Convolutional neural network architecture for RoboEM processing projections of neurite-aligned subvolumes of EM input data and making predictions about neurite continuation in terms of a curvature vector; EM data from (Motta et al., 2019). This curvature vector is representative of a parabola approximation to the neurite centerline and can be integrated during recurrent inference to yield the next position and orientation and thereby the next subvolume to be processed. Performing random rotations around the flight direction during inference can be considered zero-cost test time augmentation that allows to decorrelate subsequent inputs with respect to the orientation. The sequence of visited points during recurrent inference represents a skeleton reconstruction of the neurite. (b) Training iterations using a membrane avoiding flight policy for off-center and off-direction inputs and comparing ReLU and ELU activation functions (Clevert et al., 2015; Glorot et al., 2011), as well as EM only input versus EM and membrane probability maps as input. Membrane probabilities were found to only help in the initial training phase, while ultimately EM data is sufficient for predicting neurite continuation. Random rotation inference mode as depicted in (a) always outperforms the normal inference mode. The ELU activation function empirically performs better than ReLU. (c) Reset-based evaluation of RoboEM for different step size factors $f$ on the test set axons from the mouse cortex dataset (Motta et al., 2019). While most evaluations in this work are based on $f = 1$, the step size can be increased up to $f = 5$ yielding about 5-fold faster inference without negatively impacting the performance of RoboEM.

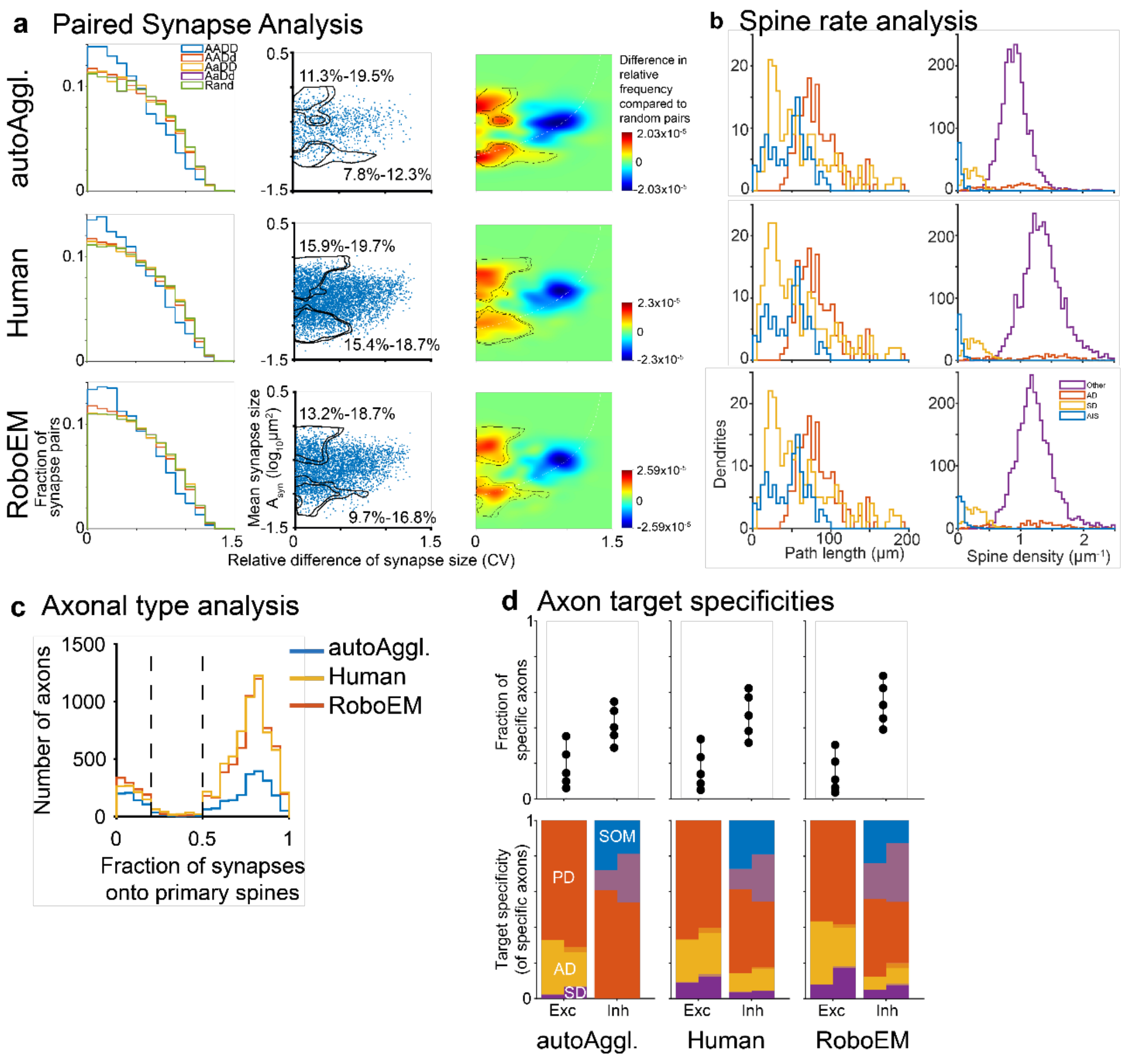


**Suppl. Figure 2. Connectomic analyses compared across reconstruction states.** Analyses based on EM data and reconstructions by (Motta et al., 2019) (a) Paired same-axon same-dendrite analysis yielding similar fractions of paired synapses that are consistent with Hebbian learning. (b) Spine rate analysis yielding underestimates of apical dendrite spine densities for the reconstruction state before RoboEM or human-based spine attachment. (c) Connectomic definition of excitatory and inhibitory axons with at least 10 synapses based on fraction of primary spine innervation. The automated state before RoboEM or human-based split and merge resolution of axons underestimates the relative proportion of excitatory and inhibitory axons, while RoboEM yields similar proportions compared to the final reconstruction state using human annotations. (d) Axonal target specificities of excitatory and inhibitory axons based on binomial null model. Human and RoboEM-based split and merge error correction allow for the detection of inhibitory axons specifically innervating apical and smooth dendrites, while this specificity is absent for the reconstruction state prior to split and merge error correction.

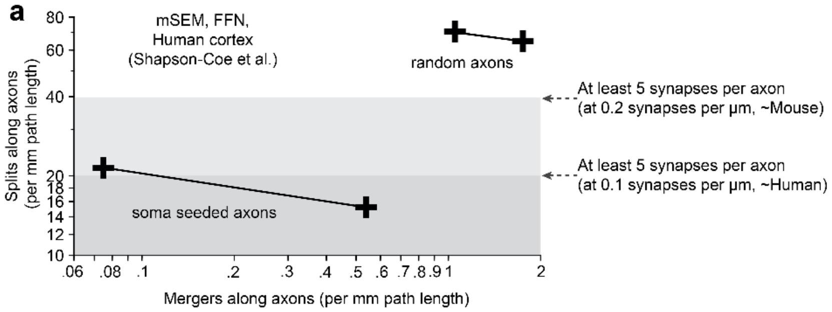


**Suppl. Figure 3. Split and merge error rates for random versus soma-seeded axons**. (a) Measured split and merge error rates for random and soma-seeded axons for the two published segmentations of FFN from (Shapson-Coe et al., 2021). Soma-seeded axons can only provide lower bounds for the reconstruction performance on random axons, which are typically thinner and therefore harder to reconstruct.
